# Exploring the potential of water channels for developing MRI reporters and sensors without the need for exogenous contrast agents

**DOI:** 10.1101/2024.01.21.576580

**Authors:** Asish N. Chacko, Austin D.C. Miller, Kaamini M. Dhanabalan, Arnab Mukherjee

## Abstract

Genetically encoded reporters for magnetic resonance imaging (MRI) offer a valuable technology for making molecular-scale measurements of biological processes within living organisms with high anatomical resolution and whole-organ coverage without relying on ionizing radiation. However, most MRI reporters rely on contrast agents, typically paramagnetic metals and metal complexes, which often need to be supplemented exogenously to create optimal contrast. To eliminate the need for contrast agents, we previously introduced aquaporin-1, a mammalian water channel, as a new reporter gene for the fully autonomous detection of genetically labeled cells using diffusion-weighted MRI. In this study, we aimed to expand the toolbox of diffusion-based genetic reporters by modulating aquaporin membrane trafficking and harnessing the evolutionary diversity of water channels across species. We identified a number of new water channels that functioned as diffusion-weighted reporter genes. In addition, we show that loss-of-function variants of yeast and human aquaporins can be leveraged to design first-in-class diffusion-based sensors for detecting the activity of a model protease within living cells.

## Introduction

Reporter genes, typically encoding fluorescent or bioluminescent proteins, are indispensable for monitoring molecular events such as gene expression, signaling pathways, and protein trafficking [1]. However, the use of optical reporters in deep tissues is hindered by the scattering and absorption of light, which impedes detailed imaging of reporter gene activity beyond a depth of approximately a millimeter from the tissue surface. To overcome this limitation, reporters whose activity can be detected using tissue penetrant imaging techniques are required. To this end, genetic reporters have been developed for several tissue-penetrant modalities [2] including photoacoustic [3], ultrasound [4,5], nuclear [6–8], and magnetic resonance imaging (MRI) [9–11]. Among these imaging methods, MRI reporters are particularly advantageous because they allow for the combination of molecular information with high anatomical resolution (∼ 100 µm in rodent models) and tomographic views of entire organs, without the use of ionizing radiation. However, most existing MRI reporters [12–19] require contrast agents, typically paramagnetic metals or metal complexes, which often need to be supplemented exogenously to create optimal contrast in reporter-expressing cells, by shortening the spin-lattice (*T*_1_) and spin-spin (*T*_2_) relaxation times of tissue water protons. This requirement to administer a synthetic (i.e., non-genetic) contrast agent is a limitation that restricts the use of MRI reporters to tissues, where the vasculature allows uniform access of cells to the injected agent.

In order to eliminate the need for adding contrast agents externally, we introduced a mammalian water channel, human aquaporin-1 (hAqp1) [20], as a new reporter gene enabling the fully autonomous detection of genetically labeled cells using diffusion-weighted MR I. This technique involves the application of pulsed magnetic field gradients separated by a user-defined interval (*Δ*) to dephase and subsequently rephase magnetization. Water molecules that diffuse to a different location within the magnetic field gradient during this interval result in a decline in net magnetization proportional to exp(-*b*.*D*), where *D* is the diffusivity of water and *b* is a parameter related to *Δ* and the strength of the gradient pulse [21]. Expression of hAqp1 facilitates the passive diffusion of water molecules across the plasma membrane. Therefore, by selecting a sufficiently long *Δ* (on the order of 100 ms [22]) to allow a substantial portion of water molecules to cross the plasma membrane, diffusion-weighted MRI can be used to visualize cells engineered to express hAqp1 based on their increased diffusivity compared to native cells (**Fig. 1a**). We and others have utilized hAqp1 to track gene expression in subcutaneous [23] and intracranial tumors [20]. Importantly, because hAqp1 does not require exogenous agents to create contrast, it has been used as a genetic reporter to noninvasively label astrocytes [24] and trace neural connectivity [25] in the mouse brain where the intact blood-brain barrier (BBB) typically hinders access to systemically injected contrast agents. A similar approach based on water exchange via the urea transporter, UT-B has also been used as a metal-free reporter gene technique to image tumor xenografts in vivo [26].

**Figure 1:**
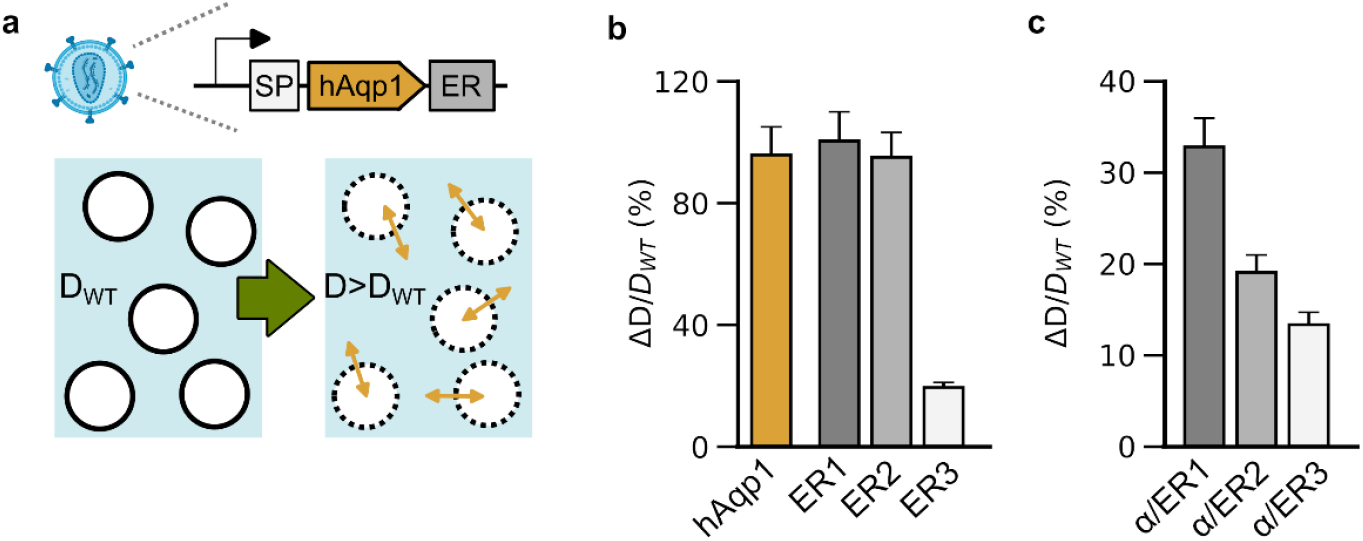
Diffusivity of cells engineered to express hAqp1 tagged with membrane trafficking motifs. **(a)** Schematic illustration of lentiviral vector depicting human aquaporin-1 (hAqp1) appended to endoplasmic reticulum (ER) export tags and an N-terminal signal peptide (SP). The diffusivity (*D*) of virally transduced HT22 cells is compared to that of wild-type cells (*D*_*WT*_) to assess the functionality of a given hAqp1 construct. **(b)** Percentage increase in diffusivity (*ΔD/D*_*WT*_) of cells expressing hAqp1 C-terminally tagged with ER export sequences derived from the glutamate receptor (ER1), Kv1.4 (ER2), and Kir2.1 (ER3). The change in the diffusivity of cells expressing hAqp1 without any export tags is shown for comparison. **(c)** Percentage increase in diffusivity of cells expressing hAqp1 with an N-terminal signal peptide derived from nAChRα in conjunction with various ER export sequences at the C-terminus. Error bars represent standard deviation from n ≥ 4 biological replicates.

Given the benefits of transmembrane water exchange as a unique mechanism for reporter gene imaging, we aimed to explore potential strategies to enhance the toolbox of MRI reporters based on water diffusion. First, we sought to determine whether modulating membrane trafficking could improve the diffusivity (and thereby, MRI contrast) of cells expressing hAqp1. We then investigated the feasibility of creating MRI contrast using water channels from diverse species to expand the range of genetic reporters available for diffusion-weighted imaging. Finally, we devised a generalizable approach to engineer functional MRI sensors via the protease-mediated regulation of water diffusivity in cells.

## Results and Discussion

### Fusing membrane trafficking motifs to hAqp1 does not lead to further enhancement of diffusivity

To explore the potential for enhancing the MRI response of aquaporins through the incorporation of motifs that regulate membrane trafficking, we designed three hAqp1 variants, each featuring distinct endoplasmic reticulum (ER) export sequences at their C-termini (**Table S1**). These ER export tags were derived from the glutamate receptor [27], voltage-gated K+ channel Kv1.4[28], and the inward rectifying K+ channel Kir2.1 [29], and have been widely utilized to enhance the cell surface expression and functionality of various membrane-localized genetic tools, including genetically encoded voltage indicators [30–32] and optogenetic tools [27,33,34]. We developed lentiviral vectors to co-express tagged hAqp1 constructs (**Fig. 1a**) with a fluorescent marker (enhanced GFP) via an internal ribosome entry site. We transduced HT22 cell lines with the above constructs and used fluorescence-activated cell sorting (FACS) to select stable cell lines based on eGFP fluorescence. Using stimulated echo diffusion-weighted MRI, as previously described in our work [22], we measured the apparent diffusivity of the ensuing cells and quantified their functionality in terms of the fold-change in diffusivity relative to wild-type cells lacking heterologous hAqp1 expression (**Fig. 1a**). Our findings revealed that cells expressing hAqp1 fused to export tags derived from Kv1.4 and glutamate receptor exhibited a similar increase in diffusivity (*ΔD/D*_*WT*_ = 95.0 ± 8.2 % and 100.4 ± 9.6 %, mean ± s.d., *P*-value < 10^−12^, n > *4*) compared to cells expressing unmodified hAqp1 (**Fig. 1b, Supplementary Fig. 1a,b**). In contrast, incorporation of the export sequence from Kir2.1 led to a significant decrease in diffusivity relative to unmodified hAqp1, producing only a moderate elevation in diffusion compared to wild-type controls (*ΔD/D*_*WT*_ = 19.5 ± 1.6 %, *P*-value = 1.8 × 10^−4^, n > *4*) (**Fig. 1b, Supplementary Fig. 1c**). Additionally, we generated cell lines expressing hAqp1 fused to a signaling peptide derived from the alpha subunit of the nicotinic acetylcholine receptor (nAChR) [27,35], in combination with each of the ER export sequences mentioned above (**Table S1**). However, cells expressing these constructs displayed a marked decrease in diffusivity compared to unmodified hAqp1 with *ΔD/D*_*WT*_ values ranging from 13.4 ± 1.4 % to 32.8 ± 3.1 % (**Fig. 1c, Supplementary Fig. 1d-f**). Collectively, these results suggest that directly appending commonly used membrane trafficking sequences to hAqp1 does not enhance whole-cell diffusion rates and may potentially interfere with endogenous trafficking motifs already present in hAqp1.

### Water channels from diverse species can function as genetically encoded MRI reporters but exhibit a lower response than hAqp1

The hAqp1 channel belongs to a group of conserved multipass membrane proteins found in diverse organisms, including plants, bacteria, and fungi [36]. Therefore, we were interested in exploring the potential of water channels from other organisms to function as genetically encoded MRI reporters. To this end, we transduced cells with lentiviral vectors engineered to express six distinct water channels (**Supplementary Fig. 2**), chosen for their high permeability coefficients: human Aqp4 [37,38] (*Homo sapiens*) and killifish Aqp0 [39] (*Fundulus heteroclitus*); plant water channels, including BvPIP2;1[40] (*Beta vulgaris*), NtPIP2;1 [41] (*Nicotiana tabacum*), and FaPIP2;1 [42] (*Fragaria* × *ananassa*); and yeast aquaporin Aqy1 from *Saccharomyces cerevisiae* (ScAqy1) [43], and *Pichia pastoris* (PpAqy1) [44]. In addition, we tested N- and C-terminal truncations of PpAqy1 and ScAqy1, specifically PpAqy1NΔ35 and ScAqy1CΔ21, which have been suggested to enhance water permeability relative to their full-length counterparts through distinct mechanisms involving pore gating [44] and membrane trafficking [43]. Using diffusion-weighted MRI, we found that hAqp4, Aqp0, and BvPIP2;1 displayed a statistically significant increase in diffusivity, ranging from a moderate elevation in BvPIP2;1 (*ΔD/D*_*o*_ = 25.1 ± 4.7 %, *P-* value = 0.03, *n* > 5) to a more substantial rise in hAqp4 (*ΔD/D*_*o*_ = 67.1 ± 11.6 %, *P-*value = 0.008, *n* = 3) (**Fig. 2a, Supplementary Fig. 3a-e**). Interestingly, full-length yeast aquaporins did not show a change in diffusivity, but their N- and C-terminal truncated isoforms exhibited a statistically significant increase, ranging from 14.9 ± 1.5 % in PpAqy1NΔ35 (*P*-value = 0.004, *n* = 6) to 55.4 ± 8.4 % for ScAqy1CΔ21 (*P*-value < 10^−4^, *n* = 6) (**Fig. 2b, Supplementary Fig. 3f-i**). Taken together, these findings demonstrate that evolutionarily distant water channels can function as diffusion-weighted reporter genes, although the fold increase in diffusivity is generally lower than that observed in cells expressing hAqp1.

**Figure 2:**
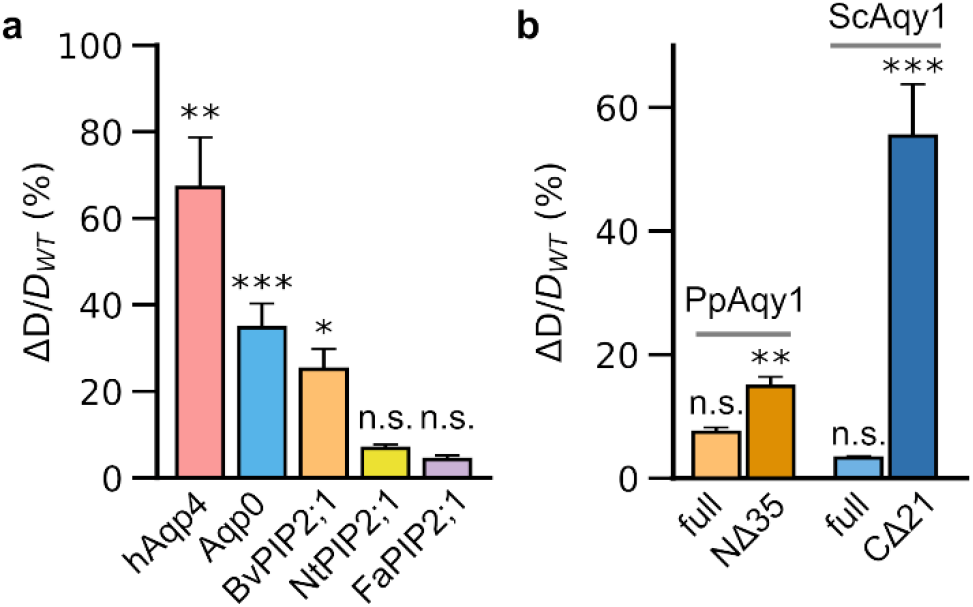
Diffusivity of cells engineered to express water channels from diverse species. **(a)** Percentage increase in diffusivity (*ΔD/D*_*WT*_) of cells transduced to express water channels from various species, including humans (hAqp4), *Fundulus heteroclitus* (MIPfun), *Beta vulgaris* (BvPIP2;1), *Nicotiana tabacum* (NtPIP2;1), and *Fragaria × ananassa* (FaPIP2;1). **(b)** Percentage increase in diffusivity of cells transduced to express full-length and truncated versions of the yeast aquaporin Aqy1 from *Pichia pastoris* (PpAqy1) and *Saccharomyces cerevisiae* (ScAqy1). NΔ35 denotes truncation of the first 35 amino acids in PpAqy1. CΔ21 denotes truncation of the last 21 amino acids in ScAqy1. Error bars represent the standard deviation from multiple biological replicates. * is *P*-value < 0.05, ** is *P*-value < 0.01, *** is *P*-value < 0.001, and n.s. is *P*-value ≥ 0.05 (Student’s t-test, 2-sided).

### ′Loss-of-function′ aquaporin variants can be harnessed to design genetically encoded sensors for detecting the activity of a model protease

Having observed that ER-tagged hAqp1 (**Fig. 1b**) and full-length yeast aquaporins (**Fig. 2b**) exhibited impaired diffusivity, we aimed to exploit these loss-of-function variants to develop genetically encoded MRI sensors of protease activity based on the proteolytic removal of N- and C-terminal sequences that interfere with aquaporin function. Our target protease was tobacco etch virus protease (TEVp), which is widely employed as a model for proof-of-concept sensor design as well as for creating programmable gene circuits to detect various biological analytes [45–47]. To create a TEVp sensor, we inserted the TEVp cleavage site into three different locations in aquaporins: following the 35-residue N-terminal pore-gating residues in PpAqy1, preceding the 21-residue C-terminal tail of ScAqy1, and sandwiched between the C-terminus and the Kir2.1 ER export sequence in hAqp1 (**Fig. 3a**). Next, we generated stable cell lines expressing each construct in com bination with doxycycline-inducible TEV protease and conducted diffusion measurements in the presence and absence of protease induction. Our results demonstrated that TEVp induction did not lead to a change in diffusivity in cells transduced with ScAqy1-TEVcs-C21 (**Supplementary Fig. 4**), indicating that protease cleavage of the C-terminal 21 residues alone was insufficient to restore function. In contrast, TEVp induction resulted in a significant increase in the diffusivities of both N35-TEVcs-PpAqy1 (ΔD/D = 33.3 ± 1.6 %, *P*-value < 10^−6^, *n* = 6) and hAqp1-TEVcs-ER^Kir2.1^ (ΔD/D = 78.4 ± 11.1 %, *P*-value = 0.001, *n* = 3) ex pressing cell lines, indicating that protease cleavage of the N-terminal capping residues and ER export motif was sufficient to restore function (**Fig. 3b, Supplementary Fig. 5**). As expected, no change in diffusivity was detected in cells expressing hAqp1 with TEVp cleavage sites incorporated within its structure [48], while maintaining unmodified N- and C-termini, indicating that TEVp-induced change in diffusion was specific to our engineered sensor constructs (**Fig. 3c**). Overall, to the best of our knowledge, N35-TEVc s-PpAqy1 and hAqp1-TEVcs-ER^Kir2.1^ represent the first-in-class MRI sensors based on the modulation of water diffusion in cells.

**Figure 3:**
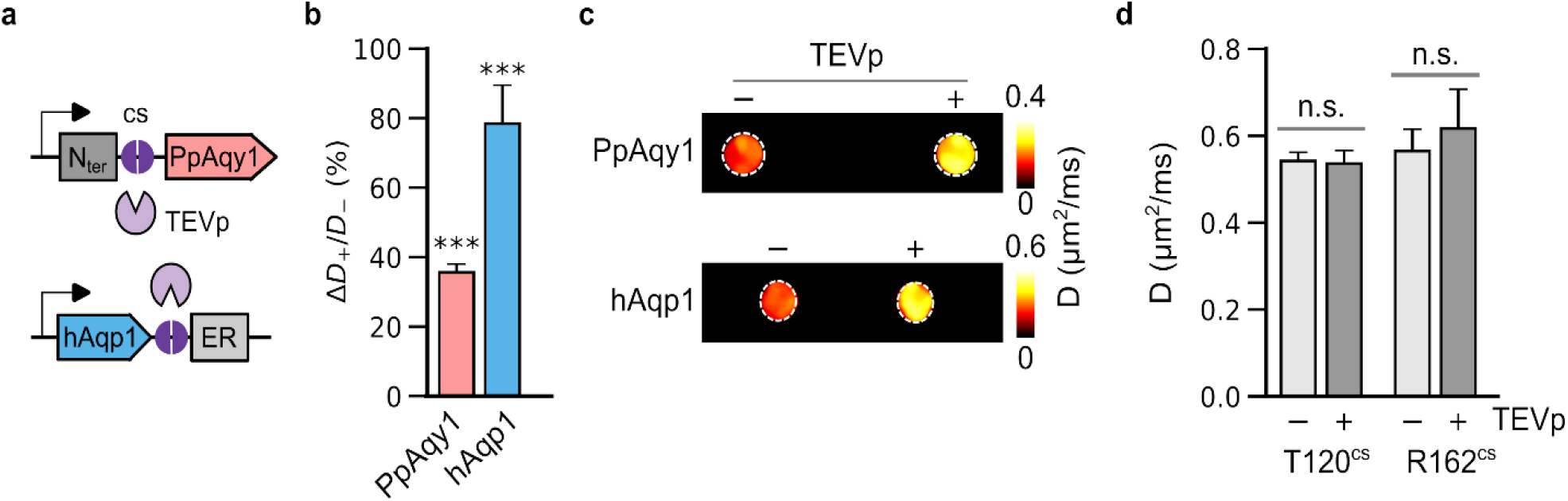
Detection of intracellular protease activity. **(a)** Schematic outline of our approach for the detection of tobacco etch virus protease (TEVp) activity based on the restoration of diffusion due to the removal of N-terminal 35 amino acids in PpAqy1 or Kir2.1 ER export sequence in hAqp1 via TEVp proteolysis at the cleavage site (cs). **(b)** Percentage increase in diffusivity (ΔD^+^/D^-^) of sensor-expressing cells in the presence of TEVp induction relative to cells in which TEVp expression was not induced. **(c)** Diffusivity maps of cell pellets with and without TEVp expression. The image is denoised using a median filter and displayed as a pseudo-colored “hotspot”. **(d)** Diffusivity of cells engineered to express hAqp1 with unmodified ends but containing TEVp cleavage sites in external (T120) or internal (R162) loop regions. The activity of TEVp is specific to the engineered sensors, as induction of TEVp expression does not produce a change in diffusivity in cells expressing hAqp1 with unmodified ends. Error bars: s.d. *** is *P*-value < 0.001, and n.s. is *P*-value ≥ 0.05 (t-test, 2-sided).

## Conclusions

In summary, we have identified a set of evolutionarily diverse water channels that can be genetically encoded in mammalian cells to produce diffusion-based MRI contrast without the need for exogenous agents. Although the newly discovered reporters generate weaker signal changes compared to hAqp1, they may serve as viable alternatives in cases where the use of hAqp1 is hindered by interference from post-translational modifications such as phosphorylation. Additionally, we found that the use of ER export sequences, which are commonly employed to improve the functionality of optogenetic tools and voltage indicators, does not further enhance diffusivity in hAqp1-expressing cells. One possibility is that membrane transit is already optimized in hAqp1, likely due to its mammalian origin. Finally, we introduced a method for engineering genetic sensors based on protease-mediated restoration of diffusion in cells expressing aquaporin variants with impaired activity. While the current study demonstrated the proof-of-concept detection of TEVp activity, we believe that this approach can be applied as a general strategy for designing sensors for a broad range of biological targets, such as protein interactions, neural activity, and signaling pathways, by combining aquaporin-driven diffusion changes with split protease technology [49,50] and synthetic gene circuits based on TEVp and related potyviral proteases [46,51,52]. As far as we are aware, only one functional MRI sensor has been genetically expressed in cells to image endogenous intracellular analytes [53], making our protease-based approach a promising solution to enhance the usefulness of MRI reporters in basic research within the field of biomedicine.

The current study has certain limitations that provide a number of opportunities for future research. First, the fairly weak response of non-mammalian aquaporins as well as some engineered variants of human Aqp1 observed in this work could be attributed to differences in membrane trafficking or the pore properties of various aquaporin isoforms. Although beyond the scope of the present study, detailed biochemical and biophysical investigations will be necessary to elucidate the precise mechanisms underlying these differences. The possibility of enhancing the performance of non-mammalian aquaporins by fusing suitable signal peptides and membrane trafficking sequences also remains to be explored. Such studies have the potential to aid in the development of improved genetic reporters as well as yield essential insights into aquaporin biology. Second, while the protease sensors engineered in this work generate a substantial change in diffusivity, there is still significant potential for optimization by increasing the on-state diffusivity. Future investigations will be required to explore ways of optimizing the on-state diffusivity, such as using appropriate spacers before the cleavage site, inserting multiple cleavage sites, and employing leucine zippers to promote protease docking at the cut site [51]. Finally, the performance and properties of these sensors in a living organism must be determined through studies in relevant animal models.

In conclusion, this study enhances the range of reporter genes that utilize transmembrane water exchange as a contrast mechanism and unveils the first generation of sensors based on this mechanism. We foresee that our research will complement progress in synthetic biology, cell-based medicine, immuno-oncology, and systems neuroscience by providing metal-free MRI reporters and functional sensors, which have long been sought as a means of measuring molecular-scale functions within intact organisms.

## Materials and Methods

### Molecular biology

ER export signals were added to the C-terminus of hAqp1 via primer overhangs during PCR amplification using a previously described hAqp1-encoding lentiviral transfer plasmid (pJY22) as the template. The tagged hAqp1 constructs were cloned into pJY22 via Gibson assembly to allow for gene expression under the control of the constitutive promoter, EF1α. Enhanced GFP (eGFP) was co-expressed via an internal ribosome entry site (IRES) to permit the identification and selection of stably transduced cells via fluorescence-activated cell sorting (FACS). The signal peptide from the nicotinic acetylcholine receptor alpha subunit (nAChRα) was obtained as a gBlock^™^ gene fragment from Integrated DNA Technologies (Coralville, IA, USA) and cloned using Gibson assembly at the N-terminus of hAqp1 directly following the start codon in pJY22. The coding regions of plant aquaporins derived from common beetroot (*Beta vulgaris*, BvPIP2;1 NCBI: AAB67870.1), tobacco (*Nicotiana tabacum*, NtPIP2;1 NCBI: NP_001313208.1), and strawberry (*Fragaria x ananassa*, FaPIP2;1 NCBI: ADJ67992.1) were obtained as gBlock^™^ gene fragments codon optimized for mammalian expression from Integrated DNA Technologies, amplified by PCR, and cloned into pJY22 using Gibson assembly. Tobacco etch virus protease (TEVp) was obtained as a gBlock^™^ and cloned into a lentiviral transfer plasmid downstream of the doxycycline-inducible minimal CMV promoter. Additionally, eBFP was co-expressed via a T2A ribosome-skipping peptide sequence to allow for the identification and selection of stably transduced cells via FACS. The TEVp sequence contained the mutations I138T, S153N, and T180A, which have been shown to improve catalytic activity [54]. All constructs were verified by whole-plasmid nanopore sequencing (Plasmidsaurus, Eugene, OR, USA) or Sanger sequencing (Genewiz, San Diego, CA, USA). All PCR amplifications were performed using the Q5® High-Fidelity 2X Master Mix (New England Biolabs, Ipswich, MA, USA).

### Construction and packaging of lentivirus

Lentiviral packaging was accomplished by combining 22 µg of a packaging plasmid (expressing viral capsid genes), 4.5 µg of envelope plasmid (encoding VSV-G to confer broad tropism), and 22 µg of transfer plasmids containing the gene of interest flanked by long terminal repeats. The plasmid mixture was brought to a final volume of 600 μL using 150 mM sterile filtered sodium chloride solution. Separately, 485 μL of linear polyethyleneimine (Polysciences, 2.58 mg/mL, 25 kDa) was mixed with 115 μL of 150 mM sterile filtered sodium chloride solution, and the contents of the two tubes were mixed by vortexing for a maximum of 10 minutes to facilitate the formation of polymeric particles incorporating all three lentiviral plasmids. Subsequently, the viral polymeric particles were added to a 10-cm plate containing HEK 293T cells (ATCC) grown to 50-70 % confluence. Approximately 16-18 h post-transfection, 10 mM sodium butyrate was added to the transfected cells to enhance the expression of the viral packaging genes. After 72 h, the supernatant was centrifuged at 500 x g for 10 min to remove cellular debris, mixed with one-tenth of the volume of Lenti-X Concentrator (Takara Bio), and incubated at 4 °C for 24 h. The lentiviral particles were then centrifuged at 1500 x g for 45 min at 4 °C and resuspended in 200 μL of DMEM and either immediately used for transduction or stored as single-use aliquots at -80 °C.

### Mammalian cell culture and engineering

HT-22, HEK 293T, and CHO Tet ON cells were cultured in a humidified incubator at 37 °C and 5 % CO_2_ in DMEM supplemented with 10 % fetal bovine serum, 100 U/mL penicillin, and 100 µg/mL streptomycin. Cells were detached using 0.25 % trypsin-EDTA for passaging. To transduce cells with concentrated lentiviral particles formed above, the cells were cultivated in a six-well plate until reaching 70-80 % confluence. After 24 h, the spent cell medium was removed from the wells, and 800 μL of a viral solution comprising concentrated lentiviral particles and 8 μg/mL polybrene was added to the cells (Thermo Fisher Scientific). Cells were then spinfected at 1048 × g for 90 min at 30 °C. After spinfection, the plates were incubated at 37 °C for 48 h. Successfully transduced cells were sorted based on eGFP fluorescence using a Sony SH800 sorter. The enriched cells were grown and stored as cryostocks until further use. TEVp-expressing cell lines were created in a manner similar to that described above and were enriched by sorting for eBFP fluorescence. These cells were subsequently re-transduced with aquaporin-encoding viral vectors and sorted for eGFP fluorescence to create doubly transduced cell lines expressing both TEVp and the TEVp-sensing constructs.

### Magnetic resonance imaging

Cells were split 24-48 h before imaging and grown to confluence in a 10-cm plate. For cells harboring an inducible TEVp construct, doxycycline hyclate (1 µg/mL) was added to the culture medium 24 hours before imaging. In preparation for MRI, the cells were detached from the plate and spun down at 350 × g for 5 min. Next, the cells were resuspended in 200 uL sterile phosphate buffered saline (PBS) and transferred to PCR tubes. The cells were spun for a second time at 500 × g for 5 min and PBS was carefully removed from above the cell pellets. Finally, the cells were resuspended in 200 µL of fresh PBS and centrifuged again to form pellets for MRI. Tubes containing cell pellets were moved to a water-filled agarose mold (1% w/v) and housed in a custom 3D-printed sample holder. All MR imaging experiments were performed using a Bruker 7 Tesla vertical-bore MRI scanner with a 66 mm diameter transceiver coil. Pellets were identified using a gradient echo localizer scan, typically with the following parameters: echo time, *T*_E_ = 3 ms, repetition time, *T*_R_ = 100 ms, flip angle, *α* = 30 °, number of averages = 1, and total acquisition time = 12.8 s. To measure diffusivity, we acquired a series of four diffusion-weighted images in the axial plane using a stimulated echo sequence with the following parameters: *T*_E_ = 18 ms, *T*_R_ = 1000 ms, gradient duration, *δ* = 5 ms, gradient separation, *Δ* = 300 ms, matrix size = 128 × 128, slice thickness = 1-2 mm, number of averages = 5, and four effective b-values in the range of 0.4-3 ms/µm^2^. Each acquisition typically required 42 minutes. All images were acquired using ParaVision software (Bruker) and analyzed using Fiji or ImageJ (National Institutes of Health). The signal intensity at a given effective b-value was computed by measuring the average intensity of all voxels inside a manually drawn region of interest (ROI) encompassing the axial view of the pellet. The apparent diffusion coefficient (*D*) was calculated as the slope of the exponential decay of the mean signal intensity as a function of the effective b-value. To generate diffusion maps, the apparent diffusion coefficient was computed for each voxel in a given ROI. The resulting image was smoothed using a median filter and pseudo-colored according to an 8-bit color scale. Least-squares regression fitting was performed by using the *fitnlm* function in Matlab (version: R2022b).

### Author Contributions

AM conceived the work. AM, ANC, ADCM, and KMD designed experiments and analyzed data. ADCM performed all experiments, including gene assembly and cloning, cell line construction, and MRI, related to the six hAqp1 constructs harboring ER export tags and signal peptides, as well as the Aqp0 construct. KMD performed all experiments (cloning, cell line construction, and MRI) involving the three plant-based water channels. ANC performed all experiments (cloning, cell line construction, and MRI) related to the two yeast aquaporins and the protease-sensing constructs. AM wrote the manuscript with inputs from all authors. All authors have read and approved the final manuscript.

## Acknowledgements

This research was supported by the National Institutes of Health (R35-GM133530, R03-DA050971, and R01-NS128278 to A.M.) and the U.S. Army Research Office via the Institute for Collaborative Biotechnologies cooperative agreement W911NF-19-D-0001-0009 (A.M.). A.D.M. acknowledges support from the Connie Frank Fellowship (2022) and The UC Regents Fellowship (2019). All MRI experiments were performed at the Materials Research Laboratory (MRL) at UC, Santa Barbara. The MRL Shared Experimental Facilities are supported by the MRSEC Program of the NSF under Award No. DMR 1720256; a member of the NSF-funded Materials Research Facilities Network. We also acknowledge the use of the Biological Nanostructures Laboratory within the California NanoSystems Institute, supported by the University of California, Santa Barbara and the University of California, Office of the President. We thank Logan Baldini for help with engineering the MIPfun-expressing cell line.

## Data availability

The data that support the findings of this study are available from the corresponding author upon reasonable request.

## Supporting Information

The following files are available free of charge: “Chacko-Miller et. al. SI 2023”. The file contains supplementary figures S1-S5, depicting multiple sequence alignment of various water channels used in this work, signal intensity versus b-value decay plots for various tagged hAqp1 constructs, and water channels sourced from diverse species, diffusion response in engineered Aqy1 from *S. cerevisiae*, and protease-dependent signal decay plots for the sensor constructs. The file also includes supplementary table S1 enumerating gene sequences of key constructs used and engineered in this work.

