## Supplementary Information for "Exploring the potential of water channels for developing MRI reporters and sensors without the need for exogenous contrast agents"

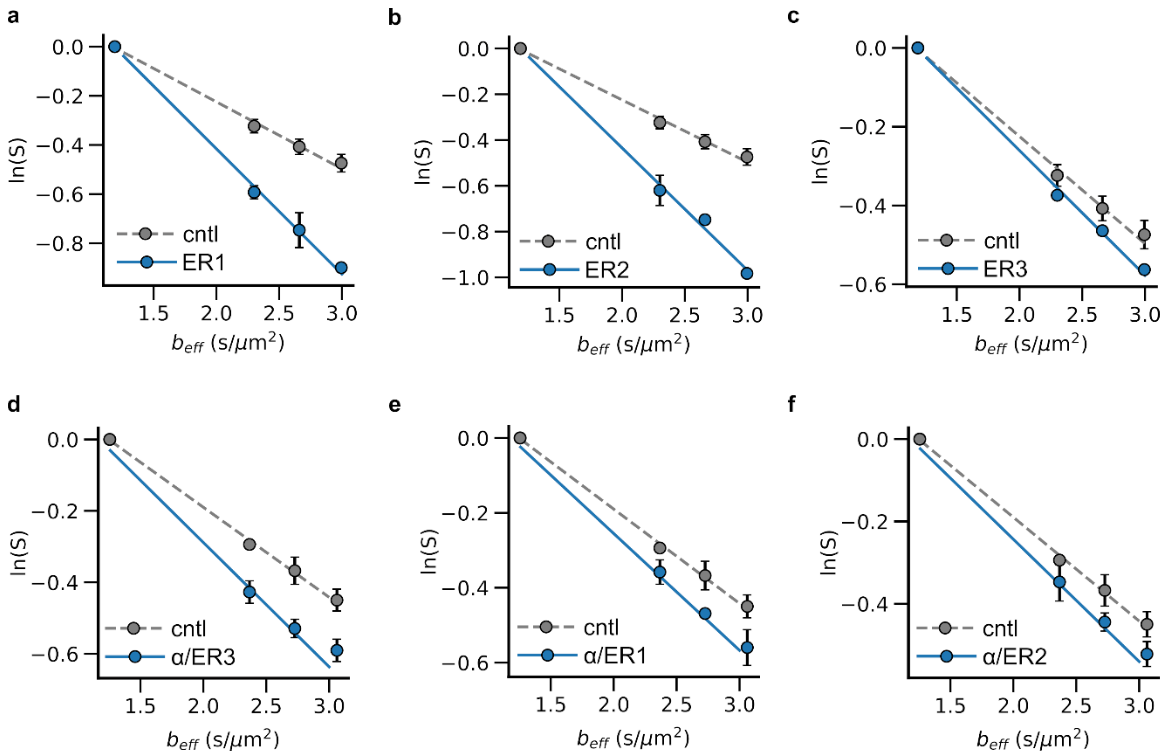


**Supplementary Figure 1**. **Diffusivity of cells expressing hAqp1 tagged with membrane trafficking motifs**. Representative plots showing the decay in signal intensity with effective b-value for HT22 cells virally transduced to express hAqp1 tagged with ER export sequences derived from **(a)** glutamate receptor **(b)** Kv1.4, and **(c)** Kir2.1. **(d)** – **(f)** show the decay plots for hAqp1 constructs harboring an N-terminal signal peptide derived from nAChRα in addition to the ER export tags. Error bars represent standard deviation from multiple biological replicates.

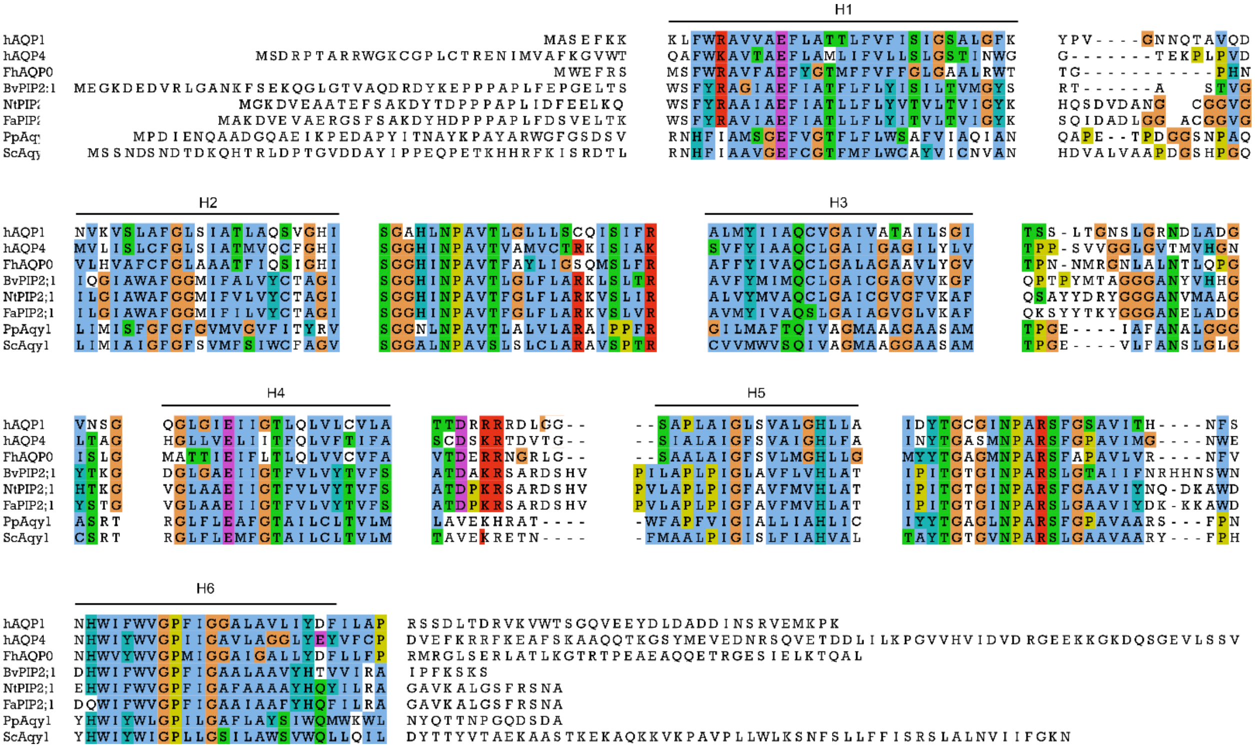

**Supplementary Figure 2**. **Multiple sequence alignment of $H_2O$ channels used in this study**. hAqp1: human aquaporin-1, FhAqp0: killifish *Fundulus heteroclitus* aquaporin-0; BvPIP2;1: *Beta vulgaris* plasma membrane intrinsic protein 2;1, NtPIP2;1: *Nicotiana tabacum* plasma membrane intrinsic protein, FaPIP2;1: *Fragaria* × *ananassa* plasma membrane intrinsic protein; Aqy1: *Pichia pastoris* aquaporin (PpAqy1) and *Saccharomyces cerevisiae* aquaporin (ScAqy1). Key conserved residues (depicted in color) are mainly localized in the membrane-spanning helices (labeled H1-H6 as per the hAqp1 structure) and in the loops connecting H2 and H3 (pseudo-helix B), and H5 and H6 (pseudo-helix E). The N- and C-termini of aquaporins are typically disordered and exhibit considerable variation in sequence composition and length across species.

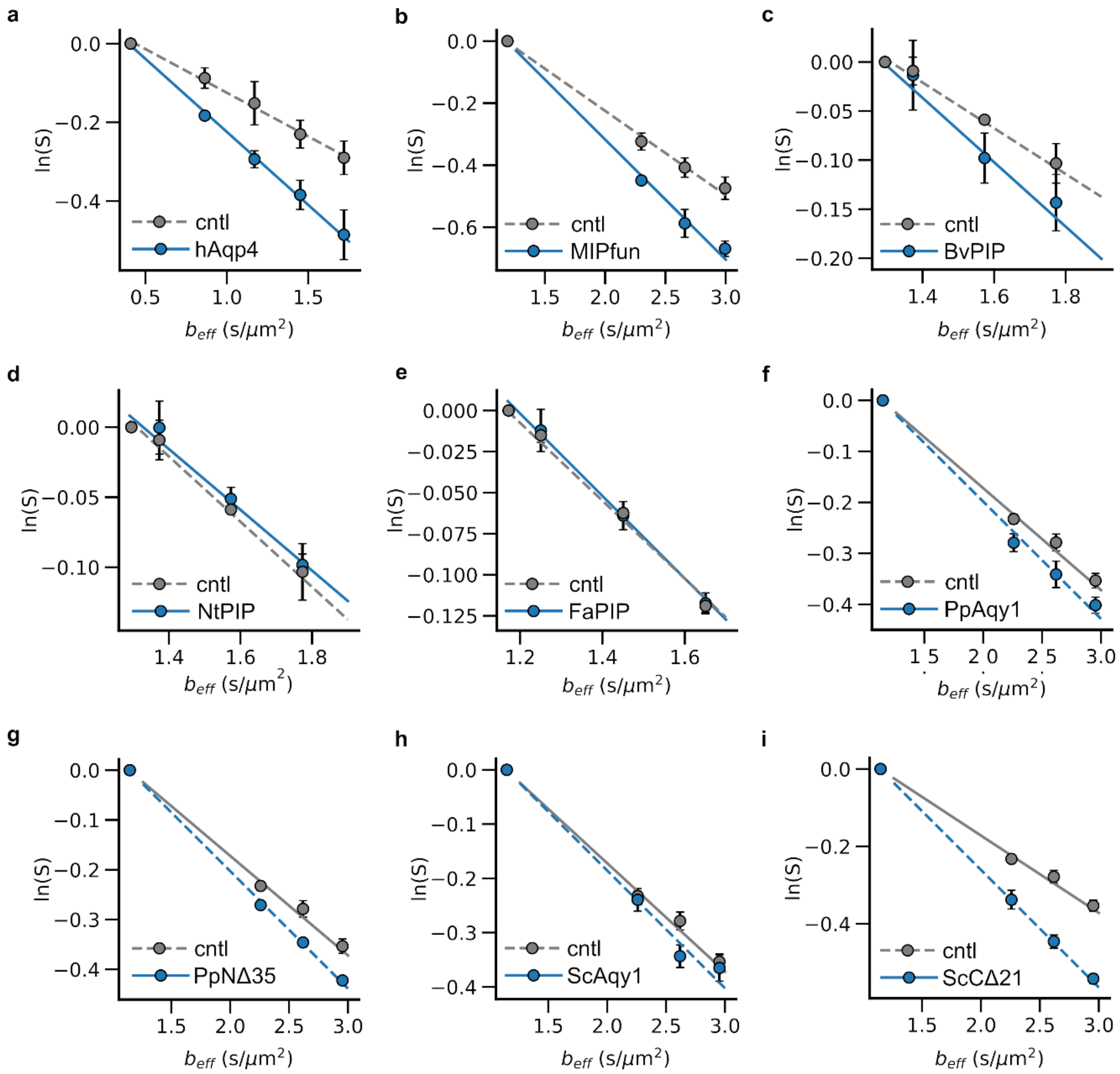


**Supplementary Figure 3**. **Diffusivity of cells expressing water channels from diverse species**. Representative plots showing the decay in signal intensity with effective b-value for CHO cells virally transduced to express animal aquaporins, including **(a)** human Aqp4 and **(b)** MIPfun/Aqp0 from the killifish *Fundulus heteroclitus*; plant water channels, including plasma membrane intrinsic proteins (PIPs) from **(c)** *Beta vulgaris* **(d)** *Nicotiana tabacum* and **(e)** *Fragaria* × *ananassa*; and fungal aquaporins, including **(f)** full length and **(g)** truncated (NΔ35) Aqy1 from *Pichia pastoris*; **(h)** full-length and **(i)** truncated (CΔ21) Aqy1 from *Saccharomyces cerevisiae*. Error bars represent standard deviation from multiple biological replicates.

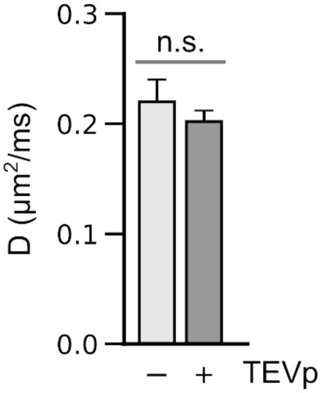


**Supplementary Figure 4**. **TEVp response of cells expressing ScAqy1-based protease sensor candidate**. Diffusivity of cells engineered to express ScAqy1 harboring a TEVp cleavage site preceding the last 21 amino acids. Error bars represent standard deviation from *n* = 6 biological replicates. n.s. denotes P-value ≥ 0.05 (Student's t-test, 2-sided).

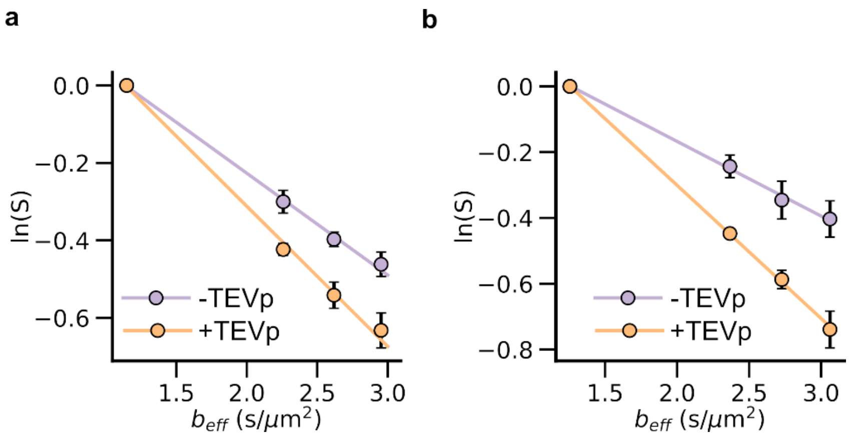


**Supplementary Figure 5**. **Diffusivity of cells expressing protease sensors based on water channels**. Representative plots showing the decay in signal intensity with effective b-values in the absence and presence of TEVp induction in CHO cells that were virally transduced to express (a) PpAqy1 harboring a TEVp cleavage sequence after the terminal 35 amino acids (b) hAqp1 containing the Kir2.1 ER export motif at the C-terminus, separated by a TEVp cleavage site. Error bars represent standard deviation from multiple biological replicates.

**Table S1. DNA sequences of water channels engineered in this work.**

| Gene | Species | Sequence |
|---|---|---|
| hAqp1 (pJY22) | *H. sapiens* | MDYKDDDDKASEFKKKLFWRAVVAEFLATTLFVFISIGSALGFKYPVGNNQTAVQDNVKVSLAFGLSIATLAQSVGHISGAHLNPAVTLGLLLSCQISIFRALMYIIAQCVGAIVATAILSGITSSLTGNSLGRNDLADGVNSGQGLGIEIIGTLQLVLCVLATTDRRRRDLGGSAPLAIGLSVALGHLLAIDYTGCGINPARSFGSAVITHNFSNHWIFWVGPFIGGALAVLIYDFILAPRSSDLTDRVKVWTSGQVEEYDLDADDINSRVEMKPK* |

| | | |
|---|---|---|
| hAqp1-ER1 (pADM11) | *H. sapiens* | MDYKDDDDKASEFKKKLFWRAVVAEFLATTLFVFISIGSALGFKYPVGNNQTAVQDNVKVSLAFGLSIATLAQSVGHISGAHLNPAVTLGLLLSCQISIFRALMYIIAQCVGAIVATAILSGITSSLTGNSLGRNDLADGVNSGQGLGIEIIGTLQLVLCVLATTDRRRRDLGGSAPLAIGLSVALGHLLAIDYTGCGINPARSFGSAVITHNFSNHWIFWVGPFIGGALAVLIYDFILAPRSSDLTDRVKVWTSGQVEEYDLDADDINSRVEMKPKRLQVMIQEAYI* |
| hAqp1-ER2 (pADM09) | *H. sapiens* | MDYKDDDDKASEFKKKLFWRAVVAEFLATTLFVFISIGSALGFKYPVGNNQTAVQDNVKVSLAFGLSIATLAQSVGHISGAHLNPAVTLGLLLSCQISIFRALMYIIAQCVGAIVATAILSGITSSLTGNSLGRNDLADGVNSGQGLGIEIIGTLQLVLCVLATTDRRRRDLGGSAPLAIGLSVALGHLLAIDYTGCGINPARSFGSAVITHNFSNHWIFWVGPFIGGALAVLIYDFILAPRSSDLTDRVKVWTSGQVEEYDLDADDINSRVEMKPKVLGSL* |
| hAqp1-ER3 (pADM10) | *H. sapiens* | MDYKDDDDKASEFKKKLFWRAVVAEFLATTLFVFISIGSALGFKYPVGNNQTAVQDNVKVSLAFGLSIATLAQSVGHISGAHLNPAVTLGLLLSCQISIFRALMYIIAQCVGAIVATAILSGITSSLTGNSLGRNDLADGVNSGQGLGIEIIGTLQLVLCVLATTDRRRRDLGGSAPLAIGLSVALGHLLAIDYTGCGINPARSFGSAVITHNFSNHWIFWVGPFIGGALAVLIYDFILAPRSSDLTDRVKVWTSGQVEEYDLDADDINSRVEMKPKFCYENEV* |
| nAChRα-hAqp1-ER1 (pADM14) | *H. sapiens* | MGLRALMLWLLAAAGLVRESLQGDYKDDDDKASEFKKKLFWRAVVAEFLATTLFVFISIGSALGFKYPVGNNQTAVQDNVKVSLAFGLSIATLAQSVGHISGAHLNPAVTLGLLLSCQISIFRALMYIIAQCVGAIVATAILSGITSSLTGNSLGRNDLADGVNSGQGLGIEIIGTLQLVLCVLATTDRRRRDLGGSAPLAIGLSVALGHLLAIDYTGCGINPARSFGSAVITHNFSNHWIFWVGPFIGGALAVLIYDFILAPRSSDLTDRVKVWTSGQVEEYDLDADDINSRVEMKPKRLQVMIQEAYI* |
| nAChRα-hAqp1-ER2 (pADM12) | *H. sapiens* | MGLRALMLWLLAAAGLVRESLQGDYKDDDDKASEFKKKLFWRAVVAEFLATTLFVFISIGSALGFKYPVGNNQTAVQDNVKVSLAFGLSIATLAQSVGHISGAHLNPAVTLGLLLSCQISIFRALMYIIAQCVGAIVATAILSGITSSLTGNSLGRNDLADGVNSGQGLGIEIIGTLQLVLCVLATTDRRRRDLGGSAPLAIGLSVALGHLLAIDYTGCGINPARSFGSAVITHNFSNHWIFWVGPFIGGALAVLIYDFILAPRSSDLTDRVKVWTSGQVEEYDLDADDINSRVEMKPKVLGSL* |
| nAChRα-hAqp1-ER3 (pADM13) | *H. sapiens* | MGLRALMLWLLAAAGLVRESLQGDYKDDDDKASEFKKKLFWRAVVAEFLATTLFVFISIGSALGFKYPVGNNQTAVQDNVKVSLAFGLSIATLAQSVGHISGAHLNPAVTLGLLLSCQISIFRALMYIIAQCVGAIVATAILSGITSSLTGNSLGRNDLADGVNSGQGLGIEIIGTLQLVLCVLATTDRRRRDLGGSAPLAIGLSVALGHLLAIDYTGCGINPARSFGSAVITHNFSNHWIFWVGPFIGGALAVLIYDFILAPRSSDLTDRVKVWTSGQVEEYDLDADDINSRVEMKPKFCYENEV* |
| hAqp4 | *H. sapiens* | MDYKDDDDKSDRPTARRWGKCGPLCTRENIMVAFKGVWTQAFWKAVTAEFLAMLIFVLLSLGSTINWGGTEKPLPVDMVLISLCFGLSIATMVQCFGHISGGHINPAVTVAMVCTRKISIAKSVFYIAAQCLGAIIGAGILYLVTPPSVVGGLGVTMVHGNLTAGHGLLVELIITFQLVFTIFASCDSKRTDVTGSIALAIGFSVAIGHLFAINYTGASMNPARSFGPAVIMGNWENHWIYWVGPIIGAVLAGGLYEYVFCPDVEFKRRFKEAFSKAAQQTKGSYMEVEDNRSQVETDDLILKPGVVHVIDVDRGEEKKGKDQSGEVLSSV* |

| | | |
|---|---|---|
| Aqp0 | *F. heteroclitus* | MWEFRSMSFWRAVFAEFYGTMFFVFFGLGAALRWTTGPHNVLHVAFCFGLAAATFIQSIGHISGGHINPAVTFAYLIGSQMSLFRAFFYIVAQCLGALAGAAVLYGVTPNNMRGNLALNTLQPGISLGMATTIEIFLTLQLVVCVFAVTDERRNGRLGSAALAIGFSVLMGHLLGMYYTGAGMNPARSFAPAVLVRNFVNHWVYWVGPMIGGAIGALLYDFLLFPRMRGLSERLATLKGTRTPEAEAQQETRGESIELKTQAL* |
| BvPIP2;1 (pKMD01) | *B. vulgaris* | MDYKDDDDKMEGKDEDVRLGANKFSEKQGLGTVAQDRDYKEPPPAPLFEPGELTSWSFYRAGIAEFIATFLFLYISILTVMGYSRTANKCSTVGIQGIAWAFGGMIFALVYCTAGISGGHINPAVTLGLFLARKLSLTRAVFYMIMQCLGAICGAGVVKGFQPTPYMTAGGGANYVHHGYTKGDGLGAEIIGTFVLVYTVFSATDAKRSARDSHVPILAPLPIGLAVFLVHLATIPITGTGINPARSLGTAIIFNRHHNSWNDHWIFWVGPFIGAALAAVYHTVVIRAIPFKSKS* |
| NtPIP2;1 (pKMD02) | *N. tabacum* | MDYKDDDDKMGKDVEAATEFSAKDYTDPPPAPLIDFEELKQWSFYRAAIAEFIATLLFLYVTVLTVIGYKHQSDVDANGDVCGGVGILGIAWAFGGMIFVLVYCTAGISGGHINPAVTFGLFLARKVSLIRALVYMVAQCLGAICGVGFVKAFQSAYYDRYGGGANVMAAGHTKGVGLAAEIIGTFVLVYTVFSATDPKRSARDSHVPVLAPLPIGFAVFMVHLATIPITGTGINPARSFGAAVIYNQDKAWDEHWIFWVGPFIGAFAAAAYHQYILRAGAVKALGSFRSNA* |
| FaPIP2;1 (pKMD03) | *F. ananassa* | MDYKDDDDKMAKDVEVAERGSFSAKDYHDPPPAPLFDSVELTKWSFYRAVIAEFIATLLFLYITVLTVIGYKSQIDADLGGDACGGVGILGIAWAFGGMIFILVYCTAGISGGHINPAVTFGLFLARKVSLVRAVMYIVAQSLGAIAGVGLVKAFQKSYYTKYGGGANELADGYSTGVGLAAEIIGTFVLVYTVFSATDPKRSARDSHVPVLAPLPIGFAVFMVHLATIPITGTGINPARSLGAAVIYDKKKAWDDQWIFWVGPFIGAAIAAFYHQFILRAGAVKALGSFRSNA* |
| ScAqy1 (pANC01) | *S. cerevisiae* | MDYKDDDDKSSNDSNDTDKQHTRLDPTGVDDAYIPPEQPETKHHRFKISRDTLRNHFIAAVGEFCGTFMFLWCAYVICNVANHDVALVAAPDGSHPGQLIMIAIGFGFSVMFSIWCFAGVSGGALNPAVSLSLCLARAVSPTRCVVMWVSQIVAGMAAGGAASAMTPGEVLFANSLGLGCSRTRGLFLEMFGTAILCLTVLMTAVEKRETNFMAALPIGISLFIAHVALTAYTGTGVNPARSLGAAVAARYFPHYHWIYWIGPLLGSILAWSVWQLLQILDYTTYVTAEKAASTKEKAQKKVKPAVPLLWLKSNFSLLFFISRSLALNVIIFGKN* |
| ScAqy1 CΔ21 (pANC02) | *S. cerevisiae* | MDYKDDDDKSSNDSNDTDKQHTRLDPTGVDDAYIPPEQPETKHHRFKISRDTLRNHFIAAVGEFCGTFMFLWCAYVICNVANHDVALVAAPDGSHPGQLIMIAIGFGFSVMFSIWCFAGVSGGALNPAVSLSLCLARAVSPTRCVVMWVSQIVAGMAAGGAASAMTPGEVLFANSLGLGCSRTRGLFLEMFGTAILCLTVLMTAVEKRETNFMAALPIGISLFIAHVALTAYTGTGVNPARSLGAAVAARYFPHYHWIYWIGPLLGSILAWSVWQLLQILDYTTYVTAEKAASTKEKAQKKVKPAVPLLWLKSN* |
| PpAqy1 (pANC04) | *P. pastoris* | MPDIENQAADGQAEIKPEDAPYITNAYKPAYARWGFGSDSVRNHFIAMSGEFVGTFLFLWSAFVIAQIANQAPETPDGGSNPAQLIMISFGFGFGVMVGVFITYRVSGGNLNPAVTLALVLARAIPPFRGILMAFTQIVAGMAAAGAASAMTPGEIAFANALGGGASRTRGLFLEAFGTAILCLTVLMLAVEKHRATWFAPFVIGIALLIAHLICIYYTGAGLNPARSFGPAVAARSFPNYHWIYWLGPILGAFLAYSIWQMWKWLNYQTTNPGQDSDADYKDDDDK* |
| Pp Aqy1 NΔ35 | *P. pastoris* | MGSDSVRNHFIAMSGEFVGTFLFLWSAFVIAQIANQAPETPDGGSNPAQLIMISFGFGFGVMVGVFITYRVSGGNLNPAVTLALVLARAIPP |

| | | |
|---|---|---|
| (pANC04) | | FRGILMAFTQIVAGMAAAGAASAMTPGEIAFANALGGGASRTRGLF<br>LEAFGTAILCLTVLMLAVEKHRATWFAPFVIGIALLIAHLICIYYT<br>GAGLNPARSFGPAVAARSFPNYHWIYWLGPILGAFLAYSIWQMWKW<br>LNYQTTNPGQDSDADYKDDDDK* |
| ScAqy1-<br>TEVcs-<br>C21<br>(pANC03) | *S. cerevisiae* | MDYKDDDDKSSNDSNDTDKQHTRLDPTGVDDAYIPPEQPETKHHRF<br>KISRDTLRNHFIAAVGEFCGTFMFLWCAYVICNVANHDVALVAAPD<br>GSHPGQLIMIAIGFGFSVMFSIWCFAGVSGGALNPAVSLSLCLARA<br>VSPTRCVVMWVSQIVAGMAAGGAASAMTPGEVLFANSLGLGCSRTR<br>GLFLEMFGTAILCLTVLMTAVEKRETNFMAALPIGISLFIAHVALT<br>AYTGTGVNPARSLGAAVAARYFPHYHWIYWIGPLLGSILAWSVWQL<br>LQILDYTTYVTAEKAASTKEKAQKKVKPAVPLLWLKSNENLYFQSF<br>SLLFFISRSLALNVIIFGKN* |
| N35-<br>TEVcs-<br>PpAqy1-<br>(pANC06) | *P. pastoris* | MPDIENQAADGQAEIKPEDAPYITNAYKPAYARWGFENLYFQ’GSD<br>SVRNHFIAMSGEFVGTFLFLWSAFVIAQIANQAPETPDGGSNPAQL<br>IMISFGFGFGVMVGVFITYRVSGGNLNPAVTLALVLARAIPPFRGI<br>LMAFTQIVAGMAAAGAASAMTPGEIAFANALGGGASRTRGLFLEAF<br>GTAILCLTVLMLAVEKHRATWFAPFVIGIALLIAHLICIYYTGAGL<br>NPARSFGPAVAARSFPNYHWIYWLGPILGAFLAYSIWQMWKWLNYQ<br>TTNPGQDSDADYKDDDDK* |
| hAqp1-<br>TEVcs-<br>ER2<br>(pANC16) | *H. sapiens* | MDYKDDDDKASEFKKKLFWRAVVAEFLATTLFVFISIGSALGFKYP<br>VGNNQTAVQDNVKVSLAFGLSIATLAQSVGHISGAHLNPAVTLGLL<br>LSCQISIFRALMYIIAQCVGAIVATAILSGITSSLTGNSLGRNDLA<br>DGVNSGQGLGIEIIGTLQLVLCVLATTDRRRRDLGGSAPLAIGLSV<br>ALGHLLAIDYTGCGINPARSFGSAVITHNFSNHWIFWVGPFIGGAL<br>AVLIYDFILAPRSSDLTDRVKVWTSGQVEEYDLDADDINSRVEMKP<br>KENLYFQ’SFCYENEV* |
| hAqp1-<br>T120$^{TEVcs}$<br>(pANC09) | *H. sapiens* | MDYKDDDDKASEFKKKLFWRAVVAEFLATTLFVFISIGSALGFKYP<br>VGNNQTAVQDNVKVSLAFGLSIATLAQSVGHISGAHLNPAVTLGLL<br>LSCQISIFRALMYIIAQCVGAIVATAILSGITSSLTENLYFQSGNS<br>LGRNDLADGVNSGQGLGIEIIGTLQLVLCVLATTDRRRRDLGGSAP<br>LAIGLSVALGHLLAIDYTGCGINPARSFGSAVITHNFSNHWIFWVG<br>PFIGGALAVLIYDFILAPRSSDLTDRVKVWTSGQVEEYDLDADDIN<br>SRVEMKPK* |
| hAqp1-<br>R162$^{TEVcs}$<br>(pANC10) | *H. sapiens* | MDYKDDDDKASEFKKKLFWRAVVAEFLATTLFVFISIGSALGFKYP<br>VGNNQTAVQDNVKVSLAFGLSIATLAQSVGHISGAHLNPAVTLGLL<br>LSCQISIFRALMYIIAQCVGAIVATAILSGITSSLTGNSLGRNDLA<br>DGVNSGQGLGIEIIGTLQLVLCVLATTDRRRRENLYFQSDLGGSAP<br>LAIGLSVALGHLLAIDYTGCGINPARSFGSAVITHNFSNHWIFWVG<br>PFIGGALAVLIYDFILAPRSSDLTDRVKVWTSGQVEEYDLDADDIN<br>SRVEMKPK* |
